# Subspecies divergence, hybridisation and the spatial environment shape phylosymbiosis in the microbiome of house mice

**DOI:** 10.1101/2023.12.11.571054

**Authors:** Susana C. M. Ferreira, Víctor Hugo Jarquín-Díaz, Aimara Planillo, Ľudovít Ďureje, Iva Martincová, Stephanie Kramer-Schadt, Sofia K. Forslund, Emanuel Heitlinger

## Abstract

Closely related host species share similar symbionts, yet how host genetics and the environment affect symbiont communities at different stages of host genetic divergence remains largely unknown. Similarly, it is unclear whether host-symbiont associations result from or contribute to host divergence.

We examined the intestinal community of 619 wild-caught mice from Germany’s European house mouse hybrid zone. Here, hybridisation upon secondary contact reflects divergence and could be traced gradually.

Temporal and spatial factors were strong predictors of microbiome composition. Subspecies divergence predicted the composition similarity of the overall microbiome, specifically in the bacteria, parasite and fungal components. The effect of hybridisation was generally weak but significant for the fungal component. We confirmed our results in experiments with wild-derived inbred mice: subspecies’ genetic distances and hybridisation predicted the overall microbiome composition, and hybridisation further predicted fungal similarities among individuals. Fungi seemed more stable to the community perturbation of infection than other components of the microbiome.

Differences between subspecies were more substantial across different microbiome components than those associated with hybridisation. Diverged microbiomes are a product of host divergence and are maintained by host genetics upon large environmental effects. These results provide a unique perspective into the ecoevolutionary processes shaping phylosymbiosis.

## Background

Microbe biodiversity is increasingly recognised as a research focus with medical and veterinary implications especially for host-symbiont associations. Similarities between host-associated microbial communities are usually congruent with the evolutionary relatedness of their respective host organisms. This observation has been conceptualised as “phylosymbiosis” [1–3]. Generally, the community of symbionts diverges at some point in the evolutionary process of host speciation. Conceptually, phylosymbiosis emerges from evolutionary processes, namely an interplay of cospeciation with potential adaptation (unilaterally by host or symbiont or even coadaptation) and ecological drift, the latter defined as the inevitable random emergence and extinction of taxa within communities [4, 5].

Observable host-symbiont communities, however, are established by this evolved filtering of symbiont communities and by ecological processes both within the host and in the host’s ambient environment (the environment outside of the host) [6]. The observable patterns of phylosymbiosis are expressed against a background of host-independent ecological processes, which are rarely addressed in the context of phylosymbiosis.

The intestinal microbiome includes taxonomically diverse bacteria, viruses, and micro- and macroeukaryotic organisms, which directly (e.g., by activating immunity) or indirectly (e.g., by metabolite production) interact with the host [7–9]. This complex community comprises host-associated residents and environmental-transient components. While bacteria dominate this ecosystem in terms of species number and biomass [10], it has become increasingly evident that other components, such as fungi and eukaryotic parasites, should not be neglected because of their profound effects on the host [11], especially on the maturation and activation of the immune system [12–14] and the overall structure of the community [15]. Despite being transient components, the host’s diet also exerts a considerable influence on the microbiome [16]. This means that interactions of the different components of the microbiome with their host can be either direct, looser via other members of the community, or nonexistent. The intimacy of interactions should impact coevolutionary processes with more directly interacting components potentially coadapting with their host. These directly associated components should be subject to host filtering. In contrast, less intimately associated components are more likely to be shaped by environmental filtering.

Phylosymbiosis has predominantly been explored within the context of reproductively isolated species, primarily from a phylogenetic perspective [17, 18]. Studies of hosts in the early stages of speciation provide a window into the potential divergence of host-associated symbiont communities. In particular, the study of intestinal communities within incipient species or populations with permeable species barriers offers a unique opportunity to decipher how the complex interplay of divergent and diverse host genetics influences the evolution and ecology of these communities [1, 19]. Hybrid zones, especially tension zones stabilised by migration and selection against hybrids [20], serve as a natural laboratory, providing insights into the influence of genetic divergence of parental species and hybrid admixture.

The house mouse hybrid zone (HMHZ) is such a tension zone. It was established at secondary contact between two subspecies of the house mouse, *Mus musculus*, which diverged in Asia 0.5 million years ago. The HMHZ in central Europe is a few hundred years old and has a width of approximately 20 km. Mice in the HMHZ are advanced (multigeneration) intercrosses between *Mus musculus musculus* (*Mmm*) and *Mus musculus domesticus* (*Mmd*) parentals at the borders of the zone [21]. The genetics of hybrids can have additive effects, meaning that phenotypes are intermediate between those of the parental subspecies. This would be visible as a linear gradient ranging from one parental phenotype to the other and is not indicative of a causal role of the trait in divergence.

Alternatively, a phenomenon called transgressive segregation can lead to more extreme trait values in hybrids compared with both parental subspecies, which would be visible in a nonlinear gradient between the two [22]. If the transgressive trait has negative fitness consequences, it might indicate causal involvement of the trait in divergence and even maintenance of species barriers [23].

Microbiome composition could be such a transgressive trait, and an aberrant microbiome in house mice might indicate dysbiosis, which could have a fitness effect [19]. In the HMHZ, a transgressive abundance of parasites (increased host resistance) has been detected [24, 25], but even for pathogens the effects on fitness are not clear [26, 27]. As fitness effects of microbiomes and the concept of dysbiosis are even more disputed [28], we here abstain from concluding effects on host health, fitness or even species barriers. In contrast to previous work [19, 29] we used the HMHZ as a model to study phylosymbiosis and the divergence of microbial communities. We contrast subspecies’ genetics and hybridisation with important ecological determinants via spatial and temporal effects.

Addressing the effects of hybridisation on species composition within a cohesive analysis is not trivial. In multigeneration admixed hybrids, admixture effects should be analysed as a gradient [24, 27]. Additionally, species composition as a response is a complex multivariate measure [30]. The analysis we present here is based on the realisation that effects on the community compositions can be tested elegantly using pairwise differences between samples as response (beta-diversity) and accounting for repeated comparisons using random effects in Bayesian multilevel models [31, 32]. These models require the predictor variables to be expressed in the same way as the response, as distances or other measures reflecting a compared pair. While this is ideal for expressing genetic distances between the subspecies, it is more complex for distances in the degree of hybridisation. Measures need to capture both shifts in the composition and changes in the variance between hybrids and parentals; we apply these in our study.

Densely sampled rodents in geographical transects of up to hundreds of kilometres provide a suitable geographical scale for investigating the effects of genetic divergence, hybridisation and separating these from ecological processes via temporal and spatial heterogeneity of the microbiomes. Spatial distances between hosts can be strong drivers of microbiome structure, in some cases even stronger than host-associated factors [33, 34]. This spatial heterogeneity can result from host social interactions that facilitate microbiome transmission [31, 35] and environmental factors that affect the host and its microbiome (environmental filtering), which are unevenly distributed across space. Additionally, spatial separation limits the dispersion of hosts and their community members, contributing to spatial patterns in the microbiome [36].

In this study, we aimed to disentangle the relative importance of subspecies’ genetic differentiation, hybridisation effects and spatial and temporal distribution in shaping the house mouse microbiome within a hybrid zone. We profiled the intestinal microbiome, including prokaryotes, micro- and macro-eukaryotes, and analysed both occurrences and abundances of the different taxa in the intestinal communities of mice captured across the HMHZ. We hypothesise that 1) subspecies differences are reflected in the intestinal microbiome composition as a product of coadaptation and host filtering and 2) that hybridisation results in genetic incompatibilities and does not allow specific coadaptation of hosts and symbionts, inducing transgressive phenotypes interpretable as aberrant microbiomes. This predicts compositional shifts and/or increased variance with hybrid admixture. 3) Spatial and temporal distances reflect several processes including environmental filtering and dispersal of microbial communities. We determined the effects of subspecies’ genetics, hybridisation, and temporal and spatial distributions in a natural population of mice. We then confirmed the subspecies’ genetics and hybridisation results in a laboratory setting with wild-derived mice from both subspecies and their first generation hybrids, before and during community perturbation by experimental infection.

## Material and Methods

This study utilises previously published datasets [37], which assessed the differential detection and quantification of *Eimeria* spp. using amplicon sequencing.

### Wild mice sampling

We trapped 672 mice using live traps on 182 farms and houses located in or close to the HMHZ between 2015 and 2018. The study area ranges from 48.09 to 53.94 degrees latitude (a 680 km long area) and from 11.46 to 14.32 degrees of longitude (a 318 km wide area) (Figure 1a). In each of five years (2015-2019), mice were trapped in September to reduce potential seasonal variation. A median of two mice per locality were captured. Mice were individually isolated in cages and euthanized by cervical dislocation within 24 h after capture (animal experiment permit No. 2347/35/2014). Individual mice were measured (body length from nose to anus), weighed, and dissected. Faecal pellets were collected from the cages in which the mice had been housed overnight and stored at −80°C after shock freezing in liquid nitrogen.

**Figure 1.**
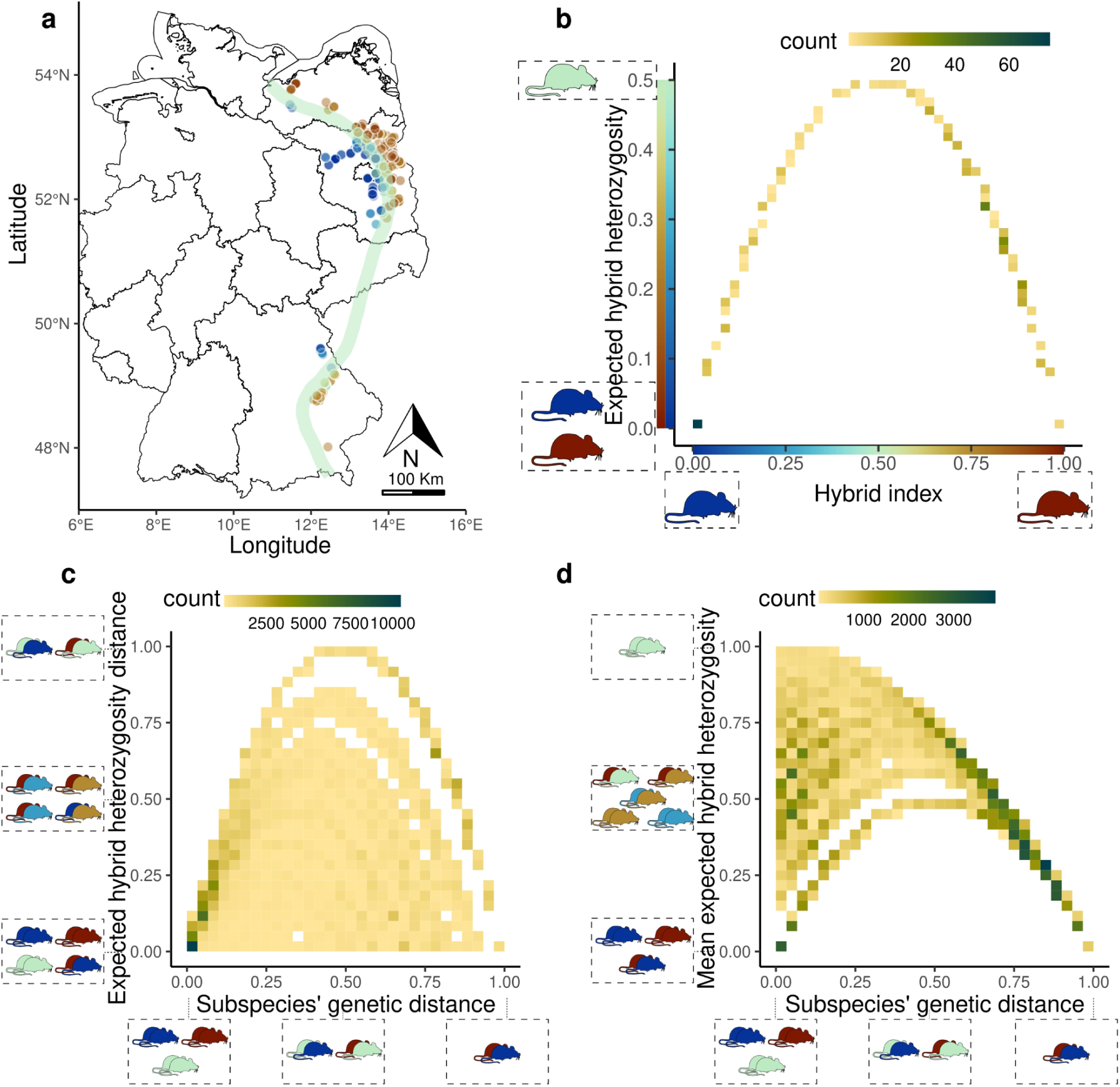
Geographical location of the study area and relationship between genetic measures derived for our analyses. **a)** The distribution of the 173 sampling locations within the house mouse hybrid zone shows 619 sampled house mice as points coloured according to a hybrid index (HI). HI was calculated as the proportion of *Mus musculus musculus* alleles genotyped at 14 subspecies-diagnostic loci. The green line represents the house mouse hybrid zone across Germany. **b)** Expected hybrid heterozygosity expresses the degree of admixture in the hybrid genotype, calculated as 2 *HI ×*(1*− HI*). Both parental genotypes have an expected hybrid heterozygosity of 0, and hybrids with equal admixture between the two subspecies have maximal values. **c)** The relationship between subspecies’ genetic distances (distance between HI values) and differences in hybridisation (hH_e_-dist) shows a strongly constrained covariance. Only when strongly admixed hybrids and parentals are compared, can high differences in hybridisation (hH_e_-dist) be assumed. Comparisons between two strongly admixed hybrids and between two pure mice (of either subspecies) have low differences in hybridisation (hH_e_-dist). **d)** Mean admixture (mean-hH_e_), which measures the average admixture in the compared pair, is similarly constrained in its covariance with subspecies’ genetic distance. Mouse pairs with large subspecies’ genetic distances are confined to pure subspecies origin and have a low mean admixture (mean-hH_e_).

### Wild-derived inbred mice sampling

Samples from four wild-derived inbred mouse strains and their F1 hybrids raised under laboratory conditions, as reported previously [38], were used to test the effect of subspecies’ genetics. In brief, parental genotypes were represented by mice from two strains: *M. m. domesticus* SCHUNT (N= 3 mice) and STRA (N= 3 mice) (Martincová et al., 2019, Piálek et al., 2008), *M. m. musculus* BUSNA (N= 3 mice) and PWD (N= 3 mice), [39, 40] and their two F1 intrasubspecies crossbreeds mus x mus (N= 3 mice), and dom x dom (N= 2 mice). In total, there were 8 mice belonging to *M. m. domesticus* and 9 mice belonging to *M. m. musculus*. Hybrids were represented by three intersubspecies-hybrid crossbreeds (N= 5 mice).

For infection, we used the Brandenburg64 isolate of *Eimeria ferrisi*, which had been isolated from the faeces of a wild *M. musculus domesticus* mouse captured in Brandenburg, Germany. We infected mice orally with 150 sporulated oocysts in 100 µl of phosphate-buffered saline (PBS 1×, pH 7.4) (experiment licence: 0431/17 issued by Landesamt für Arbeitsschutz, Verbraucherschutz und Gesundheit, Brandenburg). We collected 3–4 faecal pellets from individual mice before infection and at 6 days post infection (dpi) with *E. ferrisi*. We weighed the samples, flash froze them in liquid nitrogen and later stored them at − 80 °C for DNA extraction.

### DNA extraction

We extracted genomic DNA from faeces using the NucleoSpin®Soil kit (Macherey-Nagel GmbH & Co. KG, Düren, Germany) following the manufacturer’s protocol with the following modifications: we performed mechanical lysis of the sample in the Precellys®24 high-speed benchtop homogeniser (Bertin Technologies, Aix-en-Provence, France) using two cycles of disruption at 6000 rpm for 30 s, with a 15s delay between cycles. We eluted DNA in 40 µL TE buffer. We assessed the quality and integrity of the DNA using a full-spectrum spectrophotometer (NanoDrop 2000c; Thermo Fisher Scientific, Waltham, MA USA). We quantified the concentrations of double-stranded DNA using a Qubit® Fluorometer and the dsDNA BR (broad-range) Assay Kit (Thermo Fisher Scientific). We adjusted the DNA extracts to a final concentration of 50 ng/µl with nuclease-free water (Carl-Roth GmbH + Co. KG) and stored them at −80°C until further processing.

### Library preparation and sequencing

We used faecal DNA preparations for multimarker amplification using the microfluidics PCR system Fluidigm Access Array 48 x 48 (Fluidigm, San Francisco, California, USA). We randomised the sample order and amplified them in parallel with nontemplate negative controls using a microfluidics PCR. This allows the amplification of multiple fragments (amplicons) for different marker genes (primer pairs in additional file 1). We integrated PCR setup library preparation into the amplification procedure according to the protocol for the Access Array Barcode Library for Illumina Sequencers (single direction indexing) as described by the manufacturer (Fluidigm, San Francisco, California, USA). The amplicons were quantified using the Qubit fluorometric quantification dsDNA High Sensitivity Kit (Thermo Fisher Scientific, Walham, USA) and pooled in equimolar concentrations. The final library was purified using Agencourt AMPure XP Reagent beads (Beckman Coulter Life Sciences, Krefeld, Germany). The quality and integrity of the library were confirmed using the Agilent 2200 TapeStation with D1000 ScreenTapes (Agilent Technologies, Santa Clara, California, USA). Sequences were generated at the Berlin Center for Genomics in Biodiversity Research (BeGenDiv) on the Illumina MiSeq platform (Illumina, San Diego, California, USA) using v2 chemistry with 500 cycles. All raw sequencing data can be accessed through BioProject PRJNA548431 in the NCBI SRA.

### Identification and quality screening of the amplicon sequence variants (ASVs)

All analyses were performed in R v 4.3.1 (R Core Team, 2023). We used the R packages dada2 v. 4.3.1 [41] and MultiAmplicon v. 0.1.1 [42] to filter, sort, merge, denoise, and remove chimaeras for each run separately and for each amplicon. Additionally, we used the package decontam v. 1.21.0 [43] to remove contaminants and sequencing errors using the “prevalence” and “frequency” methods (method=”combined”). We further removed amplicon sequence variants (ASVs) that had less than 1% prevalence, less than 0.005% relative abundance [44], and samples with fewer than 100 reads. Filtering was performed individually for each amplicon in the multiamplicon datasets, and all amplicon products were collated into one “phyloseq” object using the function “merge_phyloseq” implemented in the package “phyloseq” v. 1.45.0 [45], resulting in 619 samples.

### Taxonomic annotation of the ASVs

Taxonomic assignment of the resulting ASVs was performed based on the amplicon target with the function “assignTaxonomy” from the dada2 R package [41] using the RDP classifier [46]. 18S and 16S rRNA gene sequences were classified against the SILVA 138.1 SSU Ref NR 99, and 28S rRNA gene against the SILVA 138.1 LSU Ref NR 99 databases [47], and ITS rRNA gene against the UNITE database [48]. Sequences and taxonomies from all other targeted regions that do not have a publicly available curated database were downloaded from NCBI. Sequences with more than 5 degenerated bases and lengths less than 300 bases were removed using RESCRIPt [49]. All databases were dereplicated using RESCRIPt [49].

### Merging ASVs into combined ASV (cASV)

Our dataset contained ASVs from different amplicons of different marker loci of the same taxon. Hence, ASVs were merged on the basis of their coabundance within each genus. To do so, coabundance networks for all ASVs annotated to a given genus (n=218 and n=146 for the wild and laboratory datasets, respectively) were constructed considering only positive (Pearson coefficient > 0) and significant correlations (p < 0.01) after correction for multiple testing using the Benjamini-Hochberg method. ASVs that clustered together using the “cluster_fast_greedy” function from the R “igraph” package [50] were merged into combined ASVs using the R package “phyloseq” [45], function “merge_taxa”. Here, we use the term cASVs for these combined ASVs.

Taxonomy annotation for known parasite genera was refined as developed for coccidians of the genus *Eimeria* in [37], additional file 2.

### Genotyping of mice

We genotyped mice to estimate the admixture of subspecies genomes across the HMHZ using 14 diagnostic markers. This set consists of one mitochondrial marker, one Y-linked marker, six X-linked markers, and six autosomal markers [24, 25]. Ninety-nine percent of the mice had over 10 loci amplified. For 3 of the mice, fewer loci were amplified, and the hybrid index was imputed by predictive mean matching using the R package ‘mice’ v. 3.16.0, setting meth=’pmm’ [51]. Hybridisation can result in transgressive segregation [22], making hybrids towards the centre of the HMHZ assume more extreme trait values than parentals. To create a predictor variable that captures this nonlinear effect, we calculated the expected hybrid heterozygosity (hH_e_) (Box 1). hH_e_ is the expected heterozygosity, specifically for hybrid allele combinations, and represents the degree of admixture in hybrids (Figure 1b).

##### Box 1. Definition of genetic terms

| Name | Description |
| --- | --- |
| Hybrid index (HI) | Value for each mouse calculated as the proportion of *Mmm* alleles in a set of 14 diagnostic markers. |
| Expected hybrid heterozygosity ($hH_e$) | Value ranging between 0 and 1 for each mouse that reflects the degree to which new gene combinations are brought together compared with the pure subspecies, estimated with the expected heterozygosity function: $2 HI \times (1 - HI)$. Parentals have $hH_e$ of 0 and $hH_e$ is 1 when a hybrid's genome is estimated to come equally from both subspecies [24]. |
| Subspecies' genetic distance | Pairwise difference between HI. |
| Expected hybrid heterozygosity distances ($hH_e$-dist) | Pairwise difference between $hH_e$ |
| Mean expected hybrid heterozygosity of the comparison ($hH_e$-mean) | Pairwise sum of $hH_e$. Measures how admixed the pair is as a whole. As a predictor of community differences, this variable captures whether hybrids have different variance in gut community composition. |

### Statistical modelling of community dissimilarity

Mice in our study area belong to two different subspecies or are hybrids interbred for multiple generations. Biological communities can be compared using the overall differences between the occurrence and abundance observed in each individual. To quantify the intestinal community variation (beta diversity) among mice, we calculated Jaccard distances (occurrence: presence/absence) with the function “distance” (binary = T) of the R package ‘vegan’ [52]. We repeated the analysis with Aitchison dissimilarity distances (abundance: quantitative), appropriate for use in relative abundances of taxa [53], using the function distance (pseudocount = 1). We transpose both to similarity distances (1-Jaccard distance; 1-Aitchison distance).

To test the effects of spatial and temporal distances, species barriers, and hybridisation on the β-diversity of the intestinal community, we applied Bayesian generalised linear multilevel models using the Markov chain Monte Carlo algorithm No-U-Turn Sampler (NUTS) [54] implemented in Stan through the “brms” R package v. 2.19.0 [55]. The models had intestinal community similarity as the response, and we modelled all possible pairwise distances among mice (excluding comparisons between the same individuals) as previously described [31, 32]. We used a multimembership random effects framework that allows us to account for the individuals in each pairwise comparison (e.g., Individual A, Individual B). We expressed all predictors as pairwise distances: subspecies’ genetic distance (see Box 1), expected hybrid heterozygosity distance (hHe-dist, see Box 1), mean expected hybrid heterozygosity (hHe-mean, see Box 1), spatial distance (Euclidean distances calculated from localities’s spatial coordinates), and temporal distance (difference of time a mouse pair was sampled in years). We scaled all predictors to values ranging from 0 to 1 to allow comparison of standardised estimates of the predictors. We used four Markov chains, with 4 chains, 3000 iterations, and 1000 burn-in iterations (warmup) to calibrate the Sampler, and default, uninformative priors. We visually inspected convergence and assessed the relevance of each predictor by analysing R-hat and the 95% credible intervals.

### Decomposition of the intestinal community

The intestinal community composition was decomposed into four components of cASVs: 1) bacteria including the phyla Firmicutes, Bacteroidota, Deferribacterota, Proteobacteria, Desulfobacterota, Verrucomicrobiota, Actinobacteriota, Campylobacterota, Cyanobacteria, Fusobacteriota, Patescibacteria, and unclassified bacteria; 2) fungi, including the phyla Mucoromycota, Ascomycota, and Basidiomycota; 3) plants (diet), including the phyla Anthophyta, Phragmoplastophyta, Charophyta, and Ochrophyta; and 4) parasites, including the known parasitic genera *Eimeria, Cryptosporidium, Syphacia, Aspiculuris, Mastophorus, Trichuris, Hymenolepis,* and *Tritrichomonas,* and order Ascaridida. The models were recapitulated for each component.

### Fungi and bacterial interactions

We investigated the associations between the fungal and bacteria components by modelling the bacterial composition (Jaccard and Aitchison) as a response to the fungal composition (Jaccard and Aitchison, respectively) while controlling for subspecies’ genetic distance, hHe-dist, hHe-mean, spatial distance, and temporal distance. We explored inferred interactions among 171 bacteria, 13 fungi, and 6 parasites in a co-occurrence network.

Co-occurrence networks were created with taxa with prevalence above 5% (31 samples) using the R package “SpiecEasi” [56] with the method “mb” neighbourhood selection. We used the extended method for multiple microbial domains (eukaryotes and bacteria) [57]. An optimal lambda value was observed at 0.328, and visualisation was performed using the ‘igraph’ R package [50]. Nodes with no edges were excluded from the network for visualisation.

## Results

### Intestinal community profiling in the house mouse hybrid zone

We profiled the intestinal community of 619 wild house mice, captured at 173 localities (Figure 1a), using a multimarker approach targeting both bacteria and eukaryotes [37, 58]. We derived 588 combined amplified sequence variants (cASVs; see methods) from 2880 ASVs across 34 different amplicons. We taxonomically annotated 77 genera of Eukarya and 106 genera of Bacteria and performed further analysis of the communities of cASVs.

### Strong genetic and spatial effects on overall intestinal microbiome composition in the house mouse hybrid zone

Overall, intestinal communities were more similar when mice were caught at geographically closer sites. These spatial distances were the strongest predictor for the similarities in the overall microbiome composition (Table 1, Figure 2g,h). We also observed decreased microbiome similarity in temporally distant sampling years (Table 1, Figure 2i,j).

**Table 1.**
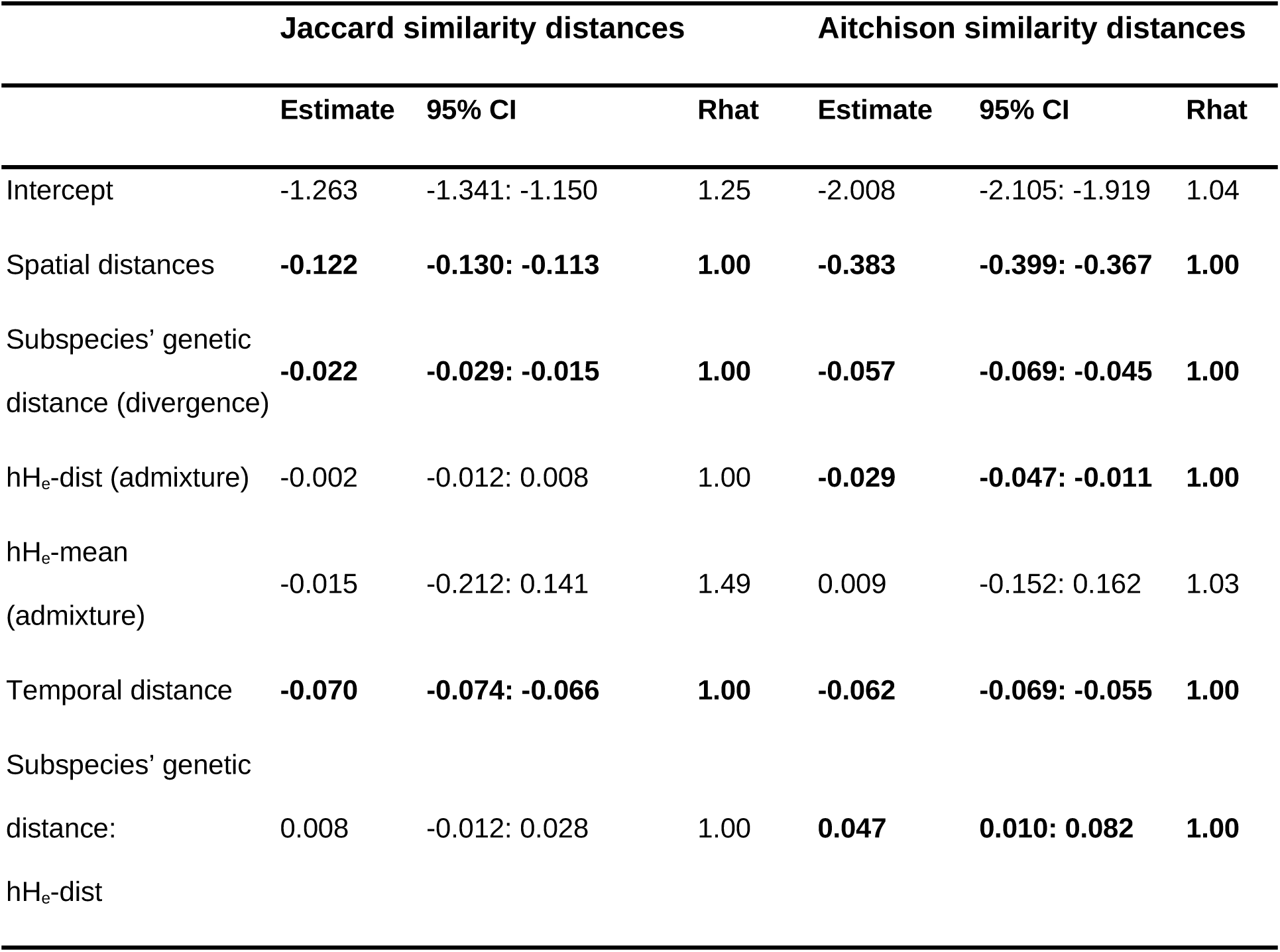
Distance-based models of the overall intestinal microbiome composition among pairs of individuals (n=191,271) for occurrence-based (Jaccard similarity distances) and abundance-based (Aitchison similarity distances). Shown are the mean estimates of the posterior distribution for each parameter and its associated 95% credible intervals (95% CI), and R-hat value that provides information on the chain convergence (if considerably greater than 1.01, the parameter estimate is not reliable). Bold highlights significant effects (credible intervals do not overlap 0).

**Figure 2.**
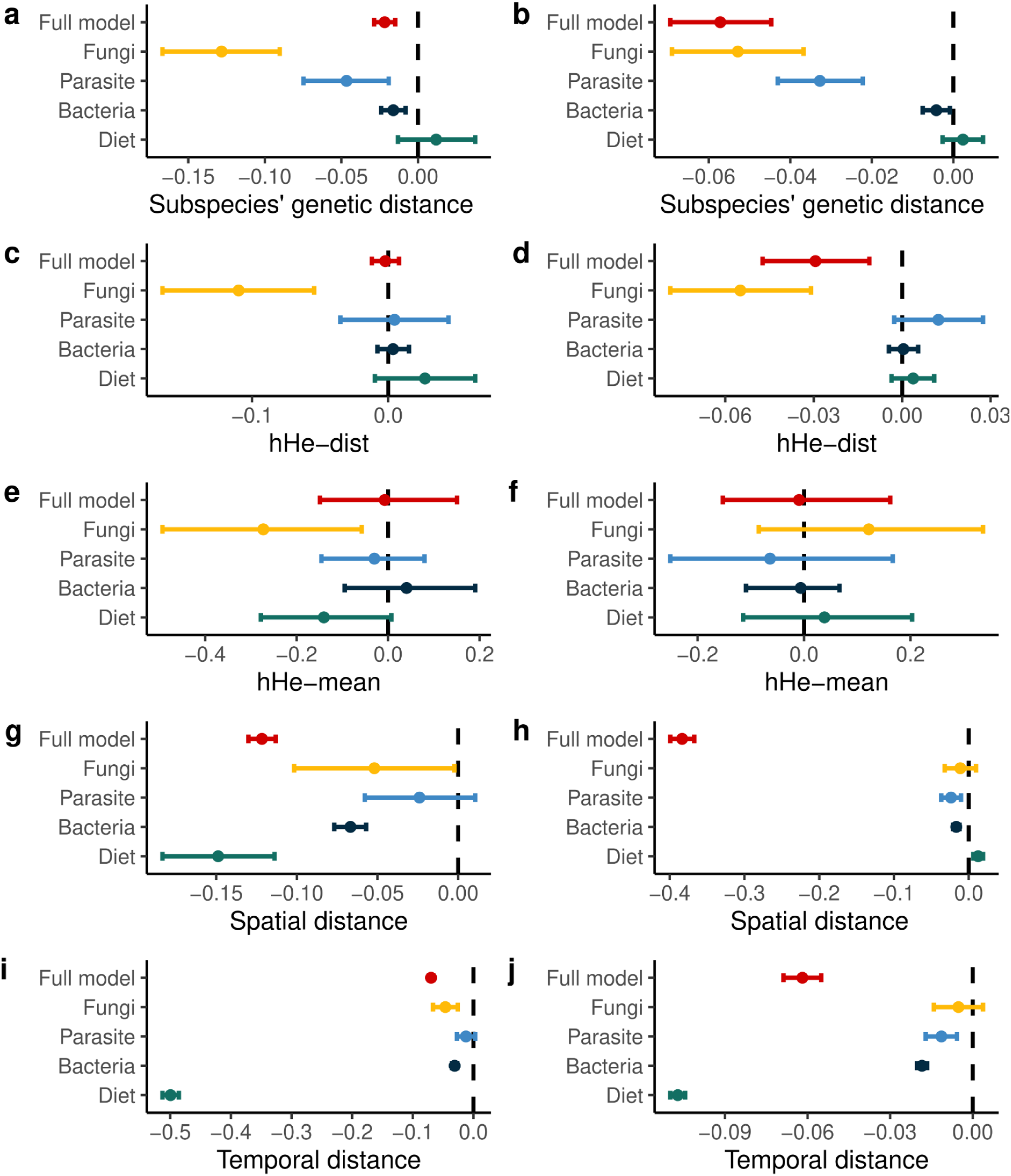
Hybridisation leads to aberrant fungal microbiomes. The compositional differences (Jaccard and Aitchison similarity distances) between whole intestinal communities (full model, 588 combined amplicon sequence variants (cASVs), red) and decomposed community components were analysed. Further colours represent the models for the community components fungi (65 cASVs, yellow), parasites (11 cASVs, light blue), bacteria (383 cASVs, dark blue) and diet (45 cASVs, green). The figure shows posterior distributions of the predictor variables: **a)** and **b)** subspecies’ genetic distances; **c)** and **d)** difference in hybridisation (hH_e_-dist); **e)** and **f)** mean admixture of the compared pairs (hH_e_-mean); g) and h) spatial distances; and **i)** and **j)** temporal distances in years. While bacteria, fungi, and parasites are significantly affected by subspecies differences, only the fungal component is affected by variables related to hybridisation. Dots represent the mean effect size, and estimates and bars represent their corresponding credible intervals (level 95%) on intestinal community composition similarity.

Most importantly, an increase in the subspecies’ genetic distances, i.e. divergence, decreased the similarities in the overall intestinal microbiome. Hybridisation had a detectable effect on the overall intestinal microbiome, as the microbiome similarities decreased with increasing hH_e_-dist, but only when abundance-based symbiont community distances (Aitchinson distances) were assessed (Table 1, Figure 2c,d). The mean admixture (hH_e_-mean) of each pair did not have significant effects on either abundance- or occurrence-based community differences (Table 1, Figure 2e,f). This means that we did not detect effects of hybridisation on the variance of community composition.

### Genetics and spatial effects on the components of the intestinal microbiome in the house mouse hybrid zone

Our approach used multiple amplicons in eukaryotic and bacterial marker genes to recover different components of the intestinal community: eukaryotic parasites, fungi, plants (diet) and bacteria. We further investigated the effects of the subspecies’ genetics, hybridisation, and spatial and temporal factors on each of the intestinal community components separately (Figure 2, Table S1, S2). Genetic distances between subspecies were associated with differences in the occurrence and abundance of parasites, fungi, and bacteria, but did not affect diet components. Differences in hybridisation (hH_e_-dist) were significantly associated with reduced fungal community similarities: mice admixed to a similar degree share more similar fungi. In addition, we found that the fungal occurrence-based composition decreased with mean admixture (hHe-mean), reflecting a slight increase in variance in the fungal community with hybridisation. A positive interaction effect of differences in hybridisation (hH_e_-dist) and subspecies’ genetic distance means that while high differences between communities are observed when hybrids are compared with pure individuals, this applies mostly to comparisons among hybrids with a similar degree of admixture. When hybrids have different subspecies’ genetics, they tend to have more dissimilar intestinal communities (Figure 3): at a subspecies’ genetic distance of 0.63, the effect of differences in hybridisation changes direction. More differently hybridised mice with a higher subspecies’ genetic distance are more similar to each other than more similarly hybridised mice with the same genetic distance. This effect becomes stronger (steeper slope) with increasing subspecies’ genetic distances. This means that differently hybridised mice have different intestinal communities, particularly fungal communities. The number of between-mouse comparisons, however, is larger (133,499 comparisons) below than above a subspecies’ genetic distance of 0.63 (57,772 comparisons). We found no evidence of the effect of mean admixture (mean-hH_e_), accounting for increased variance and hence “aberrant microbiomes” in hybrid mice, except in the occurrence-based fungi composition (Jaccard similarity distances).

**Figure 3.**
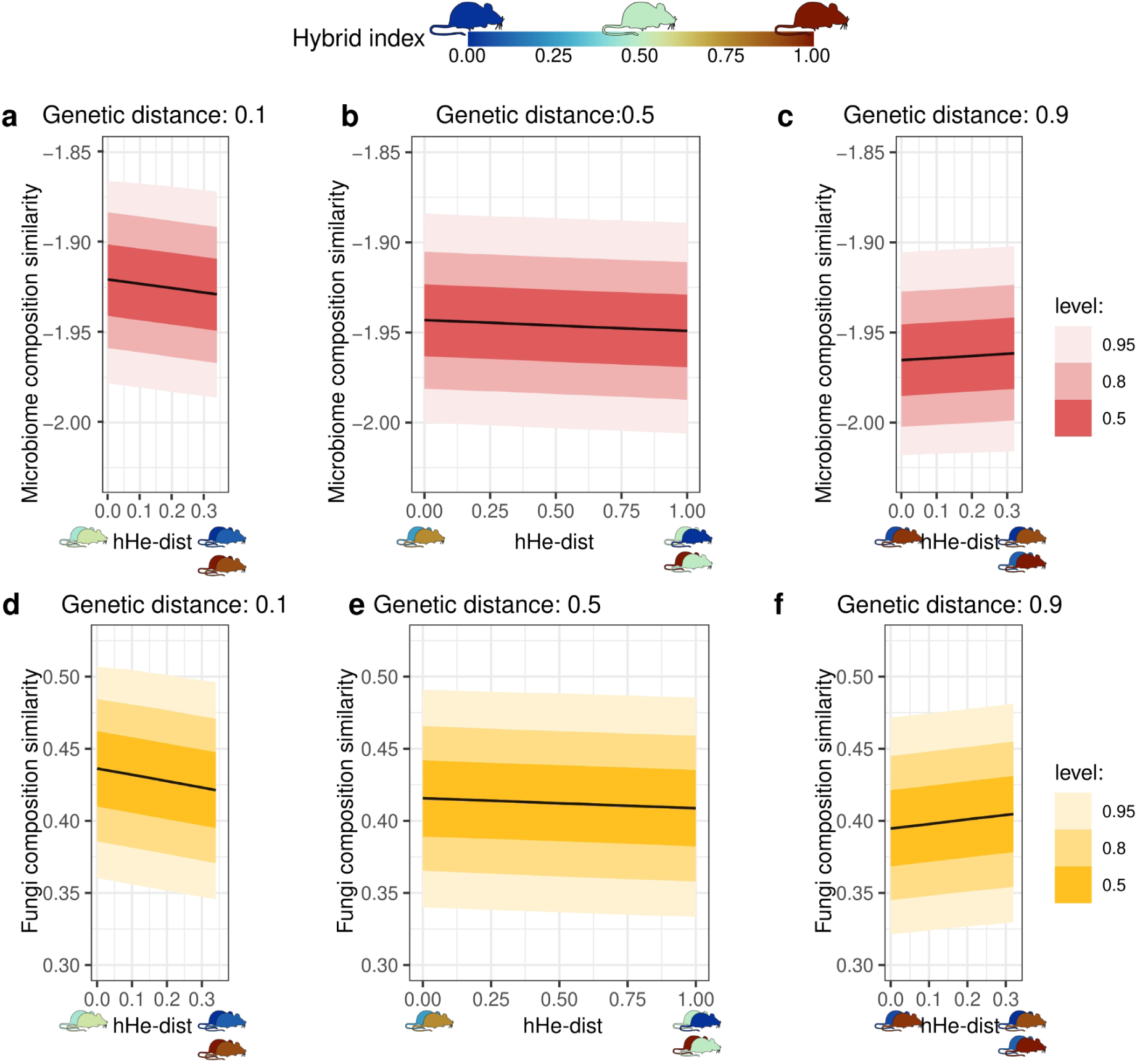
Significant statistical interaction effect between subspecies’ genetic distance and differences in hybridisation (hH_e_-dist) for the overall intestinal microbiome and fungal component compositions. **a)** At low and **b)** intermediate subspecies’ genetic distances, differences in hybridisation (hH_e_-dist) have a negative effect on the microbiome similarities. Differently hybridised mice with **d)** similar subspecies’ genetics and **e)** intermediate subspecies’ genetics show more divergent fungi composition than similarly hybridised mice. At high subspecies’ genetic distances, mice (with a large contribution from either subspecies) show higher similarity in their **c)** overall intestinal microbiome composition and especially **f)** fungal composition with differences in hybridisation. Overall intestinal microbiome and fungal composition distances are based on abundance-base composition distances (Aitchison similarity distances). The coloured mouse symbols highlight that our model is based on distances between pairs of mice (distances in the microbiome composition explained by distances of predictor variables). hH_e_-dist represents the differences in the degree of hybridisation between two mice.

The intestinal community similarity decreases with spatial distances for bacteria in both occurrence-based and abundance based compositions. Fungi and diet-derived occurrence-based and parasite abundance-based compositions decrease in similarity with increased spatial distances (Figure 2g, h). Increased temporal distances decreased the similarity of bacteria and diet composition and fungi occurrence- and parasite abundance-based compositions (Figure 2i, j).

### Fungi composition predicts bacterial composition in the house mouse hybrid zone

Effects on different components of the intestinal community might not be independent of each other; thus, we tested whether differences in the fungal community predict differences in the bacterial community. We did so while controlling for subspecies’ genetic, temporal, and spatial effects and found that the fungal community composition predicted bacterial composition in both occurrence- and abundance-based measures (Figure 4a,b, Table S3). To explore associations between taxa, we used a co-occurrence network and found 239 significant associations (edges) within 127 taxa (nodes) (Figure 4c). We found only two associations between bacteria and fungi, as the bacteria *Erwinia* and *Planococcus* were associated with the fungi *Kazachstania* and *Blumeria* and one edge between the parasite *Cryptosporidium* and the fungi *Kazachstania,* and a member of the family Saccharomycetales.

**Figure 4.**
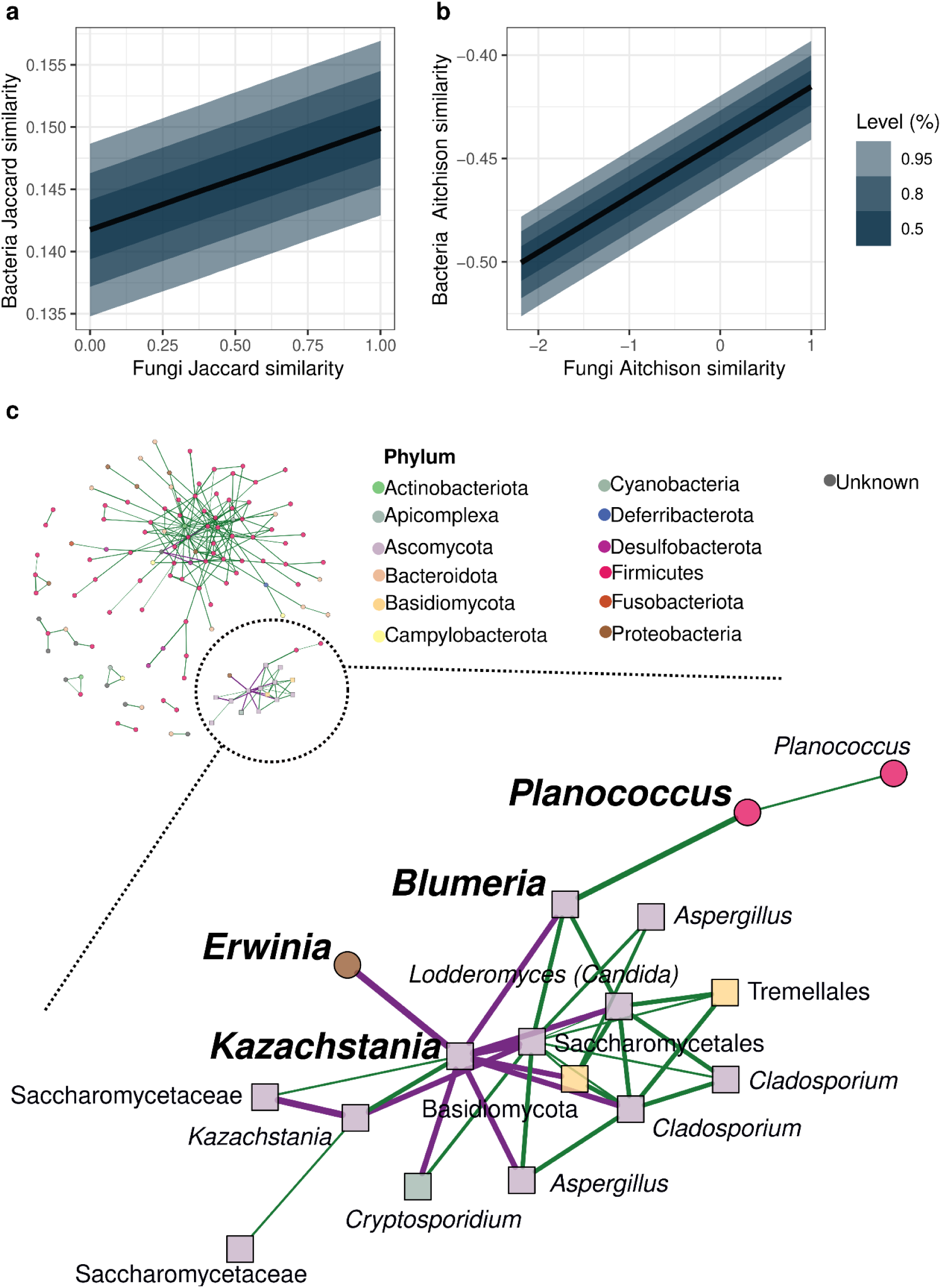
The similarity of the bacterial community increases with fungal community similarity. Independent of subspecies’ genetic, temporal, and spatial effects, mice with similar fungal communities harbor more similar bacterial communities. This is found for similarity measured both **a)** occurrence-based (which taxa are in the composition; Jaccard similarity distances) and **b)** abundance-based (at which intensity are taxa in the community found; Aitchison similarity distances). **c)** Co-occurrence network of bacteria, eukaryotic parasites, and fungi with a prevalence of at least 5% (31 samples). The segment of the network containing fungi-bacteria interactions was enlarged. Nodes correspond with combined amplicon sequence variants (cASVs), and colours correspond to the respective phyla. Circles represent bacteria and squares represent eukaryotes. Labels are the lowest taxonomic annotation for each cASV. Interacting fungi and bacteria are in bold. Edges are predicted interactions; green edges are positive and purple edges are negative. Association strength is represented by the edge thickness.

### Subspecies and hybridisation affect the intestinal community of mice in laboratory conditions before and during community perturbation by experimental infection

We profiled 22 ‘wild-derived’ inbred house mice before and at the peak of *Eimeria* infection (day 0 and day 6, respectively) in a laboratory set-up using the same multimarker approach. We derived 318 cASVs from 597 ASVs across 12 amplicons. We taxonomically annotated 36 genera of Eukarya and 81 genera of Bacteria and performed further analysis of the communities of cASVs as for the wild mice (Methods). We confirmed our findings derived from the wild mice: 1) the similarity of the overall intestinal community decreased when the subspecies’ genetic distances increased (Figure 5a, b, Table S4), and 2) the fungi occurrence-based composition and the parasite abundance-based composition decreased when the differences in hybridisation (hHe-dist) of the compared pairs increased (Figure 5c, d, Table S3).

**Figure 5.**
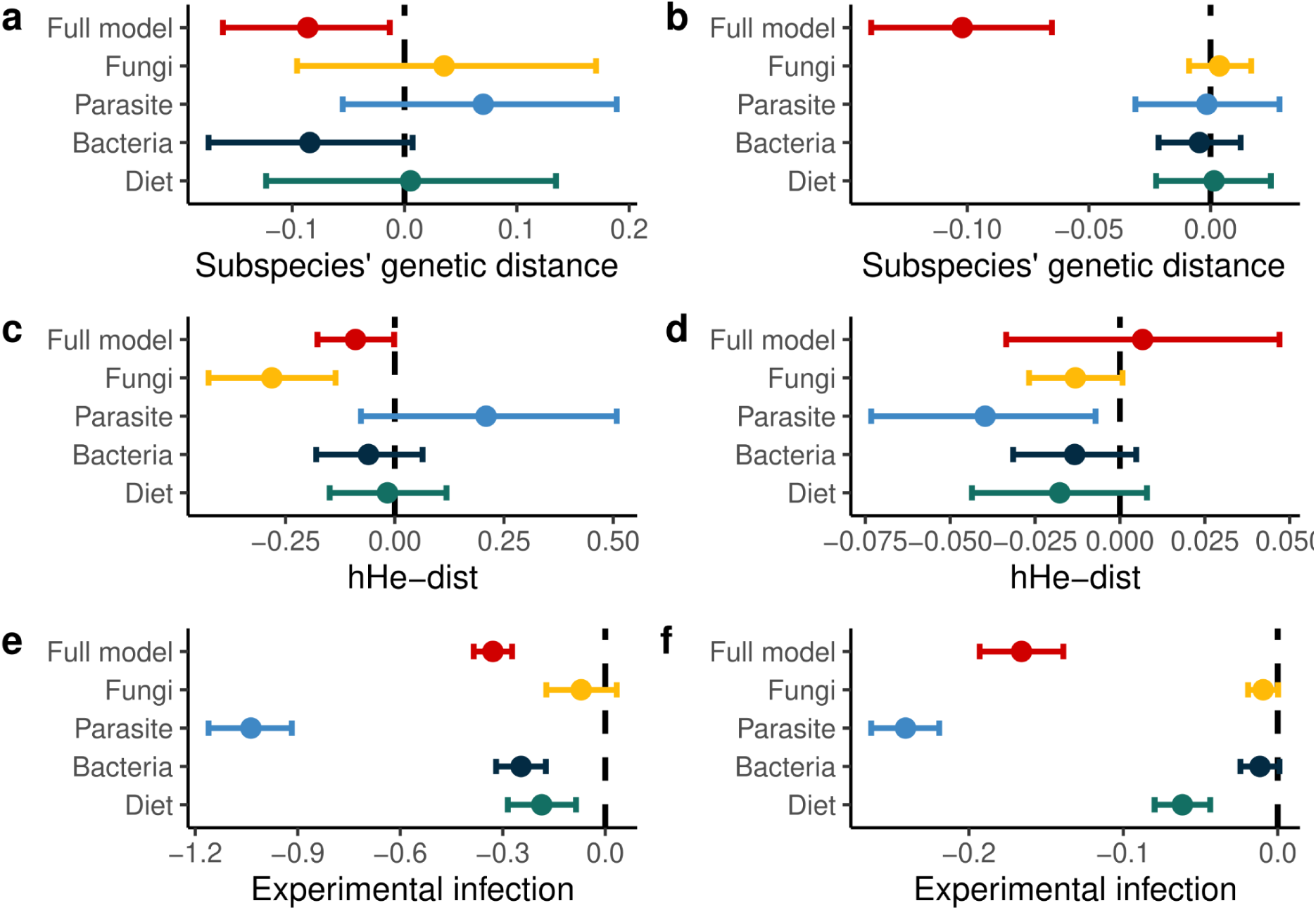
The fungal community of wild-derived inbred mice is shaped by hybridisation effects. The compositional differences (Jaccard and Aitchison similarity distances) between whole intestinal communities (full model, 318 combined amplicon sequence variants (cASVs), red) and decomposed community components were analysed for 22 mice before *Eimeria ferrisi* infection (day 0) and at the peak of infection (day 6). Further colours represent the models for the community components fungi (29 cASVs, yellow), parasites (6 cASVs, light blue), bacteria (207 cASVs, dark blue) and plants (30 cASVs, green). The figure shows posterior distributions of the predictor variables: **a)** and **b)** subspecies’ genetic distances; **c)** and **d)** differences in hybridisation (hH_e_-dist); **e)** and **f)** community perturbation (experimental infection with *Eimeria ferrisi*). Only the overall intestinal microbiome is significantly affected by subspecies differences, whereas only the fungal component is affected by hybridisation. Dots represent the mean effect size estimates and bars represent their corresponding credible intervals (level 95%) on intestinal community composition similarity. The model also accounted for infection status. While *Eimeria* infection shows an overall perturbation of the intestinal microbiome, this was not reflected in the fungal component, specifically (Table S4).

## Discussion

Phylosymbiosis, the reflection of evolutionary history in the similarity of microbiomes in related hosts, means that differences in the intestinal community are established at some point in the speciation process [59, 60]. In mice, both divergence between subspecies and hybridisation have been reported to affect the bacterial microbiome [19] and specific eukaryotic parasites [24, 25, 27]. We found that genetic differentiation between the two subspecies of the house mouse has a strong impact on the intestinal community with regard to both the occurrence and abundance of taxa. We show this in natural populations accounting for spatial and temporal distances in our sampling, which complements very recent independent work [29]. We also extended this in a laboratory setting, showing that subspecies’ genetic differentiation was also evident in wild-derived inbred mice before and after infection with *Eimeria ferrisi,* simulating a natural community perturbation. The overall effect of hybridisation was small but significant for the fungal component of the community: hybrids had a fungal composition distinct from parental mice, both in natural populations and wild-derived inbred mice. These results suggest that intestinal communities are unlikely to reinforce reproductive isolation between incipient species but point to host filtering; i.e. selection of taxa by the host internal environment. Our study provides a novel window into genetic vs. environmental effects on the microbiome and symbiont communities at large.

First, we showed that genetic differences between subspecies of the house mouse are associated with differences in the intestinal microbial community in both natural populations and in a laboratory setting. This indicates species-specific filtering of the intestinal community by host processes, independent of spatial and temporal effects. It is important to note that this present-day association of specific intestinal communities with their hosts does not necessarily indicate coadaptation or even a role of these communities in the speciation process. During the early divergence of house mouse subspecies ecological drift might very well have been the predominant force of community divergence. The close proximity of mice from different subspecies in the HMHZ, large distances between mice of the same subspecies and our spatially explicit model, however, mean that subspecies differences observed now are supported by filtering. Filtering by the host, within the intestinal environment, might be mediated by genotype-specific immune responses. Such host filtering then leads to species sorting, i.e. promotion or suppression of taxa in the community [61]. As an alternative, host characteristics such as behaviour [62, 63] could differentiate the abiotic and biotic ambient environment between the two different host subspecies. Interspecific interactions within the community could then extend the effects of symbiont colonisation and persistence [7, 64, 65]. This means that even subspecies’ genetic differences might select the microbiome via environmental filtering (in the ambient environment of the host). We addressed this in the laboratory, showing that wild-derived inbred mice had species-specific microbiome compositions and that subspecies’ genetics filter the microbiome via host physiology (e.g. the immune system) rather than the environment.

Hybridisation effects can point to species barriers, as genetic incompatibilities might induce transgressive traits that negatively affect fitness in hybrids [66]. A previous study investigating the bacterial microbiome in mice showed a differentiation between hybrids and pure subspecies in wild-derived strain representatives and their second-generation (F2) in a laboratory setting and in wild populations, and reported extreme abundance patterns linked to immune genes for specific microbial taxa such as *Helicobacter* or *Blautia* in *Mmm* [19].

Such patterns or so-called transgressive phenotypes in taxa abundances of the microbiome have a genetic basis in nonadditive allelic effects within a locus (overdominance or underdominance) or between loci (epistasis) [22]. While we cannot confirm this effect in the bacterial microbiome, we find, similar to a very recent study [29], that changes in the fungal microbiome are weakly associated with hybridisation. We also detected a small increase in the variance of the occurrence of fungi, meaning that aberrant fungal microbiome compositions are found among pairs of hybrids within natural populations. The hybrid first-generation (F1) of wild-derived inbred mice also had a fungal microbiome composition distinct from that of parentals (pure mice) in the laboratory. Hybrids are a “genetic accident” and do not form a hybrid population that can coadapt with symbionts. This can be understood in both geographical dimensions and with respect to adaptation in a hybrid tension zone: first, the zone is narrow and parentals are spatially close to hybrids [21].

Second, parentals have higher fitness than hybrids, and any hybrid should prefer a parental for mating over another hybrid. This is called “reinforcement selection” and is a factor in keeping the zone stable [67]. Taking into account the independent evidence from two studies in wild populations and in the laboratory, we can argue that fungal symbionts might be affected by hybridisation. Interestingly, the fungal component is also less affected by environmental filtering (spatial distances) in the wild and not affected by community perturbation during an experimental infection under laboratory conditions. This could point to a weaker effect of the environment (and generally ecology) on fungi than on other microbiome components. Given the tight niche-based interactions between the intestinal environment and its microbiome, hybridisation might impose a significant ecosystem filtering through intestinal dysregulation (e.g. potentially inflammation). These conditions initiate ecological filtering and selection for specific microbial populations that are metabolically independent and resilient to such conditions and can self-sustain in the absence of ecological services associated with a homeostatic environment [68]. Our results suggest a less resilient capacity of the fungal component to the conditions imposed by hybridisation but not by infection of a host-specific parasite, which is highly prevalent in natural populations. Taken together, our results suggest that fungal symbionts might be involved in earlier stages of speciation than other intestinal components. Bacteria-fungi interactions are widespread in communities, including intestinal communities [15, 69]. The fungal component is a primary source of secondary metabolites (e.g. natural antimicrobial compounds) that could affect the presence and abundance of bacteria and shape microbial communities at taxon level and thus the gene content (e.g. virulence factors, ARGs) [13]. We found that pairs with similar fungal compositions also had similar bacterial compositions, however, we detected very few individual associations of abundances between fungi and bacteria. Collectively, our results suggest broader community interactions among bacteria and fungi, e.g., through host-filtering. Transmission, colonisation and host-control of the fungal component of the microbiome are a frontier for further research both in the laboratory and in the natural environment.

## Conclusion

We differentiate subspecies’ genetic effects, indicating the filtering of symbionts by the host, from spatial-temporal effects pointing to environmental filtering. Our finding of different intestinal symbiont communities in the two house mouse subspecies suggests present-day subspecies’ genetic differences between host-symbiont interactions with unknown fitness consequences. Present-day aberrant effects of hybridisation on symbionts could suggest the involvement of symbiont interactions earlier in the process of speciation. However, we found only weak signals for an aberrant fungal component of the microbiome in hybrids. In the wild, filtering by hosts (e.g. immune regulation and control of susceptibility) cannot be completely distinguished from indirect filtering by the ambient environment (environmental filtering due to slight divergence of the species’ environmental niche). We addressed this in an experiment, including a perturbation of the community. Our laboratory results suggest direct host-mediated filtering of the microbiome and stability of hybrid-differences, especially in fungi against community perturbation. Future studies should combine immune measures and higher resolution of environmental variables (e.g. microclimate, agricultural practices or general land-use) to disentangle the different contributions, coupled with experimental set-ups of wild-derived parentals and hybrids. A focus of this research should be on fungi, while interactions between taxa in the community also need attention.

## Availability of data and materials

The datasets supporting the conclusions of this article are available in the Github repository, https://github.com/ferreira-scm/Hybridization_spatial_HMHZ. All sequencing raw data can be accessed through the BioProject PRJNA548431 in the NCBI SRA.

## Acknowledgements

We acknowledge the help of several generations of beautifully diverse undergraduate students during our sampling excursions. Their dedication to research, overhour work and youthful recklessness venturing in the wilderness of rural Germany were essential to this project. We acknowledge the open Access Publication Fund of Humboldt University Berlin.

## Abbreviations

ASV: amplicon sequence variants
cASV: combined amplicon sequence variants
hH_e_: Expected hybrid heterozygosity
hH_e_-dist: Expected hybrid heterozygosity distances
hH_e_-mean: Mean expected hybrid heterozygosity of the comparison
HI: Hybrid index
HMHZ: house mouse hybrid zone
*Mmm*: Mus musculus musculus
*Mmd*: Mus musculus domesticus

## Funding

Open Access funding enabled and organised by Projekt DEAL. Open Access Publication Fund of Humboldt University Berlin. This work was supported by the Deutsche Forschungsgemeinschaft (DFG) (Grant Number: 285969495/HE 7320/2–1; Grant Number: MA 9054/1-1 [SCMF]), the German Academic Exchange Service (DAAD) (VHJD scholarship holder during PhD studies) and the Research Training Group 2046 “Parasite Infections: From Experimental Models to Natural Systems” (VHJD associated PhD student/EH Senior Researcher). VHJD is funded by the Deutsche Forschungsgemeinschaft (DFG) grant Integrative evolutionäre und ökologische Analyse von Antibiotikaresistenzen: Auftreten und Verbreitung vom bakteriellen Genom bis zur geographischen Landschaft FO 1279/6-1 | HE 7320/5-1 | KR 4266/4-1. VHJD and SKF are supported through the JPIAMR-EMBARK project funded by the Bundesministerium für Bildung und Forschung under grant number F01KI1909A.

## Author information

### Contributions

SCMF, VHJD and EH designed the study. VHJD, EH, LD and IM collected the samples. VHJD performed laboratory and molecular work. SCMF performed the analysis. SCMF, VHJD and EH wrote the manuscript, with contributions and feedback from AP, SKF and SKS. All the authors have read and approved the final manuscript.

### Corresponding author

Correspondence to Susana C. M. Ferreira and Víctor Hugo Jarquín-Díaz

## Ethics declarations

### Ethics approval and consent to participate

Experiment licence 0431/17 issued by Landesamt für Arbeitsschutz, Verbraucherschutz und Gesundheit, Brandenburg. On the basis of §8 of the Animal Protection Act Tierschutzgesetzes), State Office for Health and Social Affairs–Berlin (Landesamt für Gesundheit und Soziales–Berlin [LAGeSo]) hereby grant the permission to Prof. Dr. Emanuel Heitlinger to carry out scientific experiments on live vertebrates; a total of 1080 mice *Mus musculus*. Permit to capture house mice: capture permit No. 2347/35/2014.

### Consent for publication

Not applicable.

### Competing interests

The authors declare no competing interests.

## Supplementary information

**Additional file 1:** Spreadsheet containing description and references of the primer pairs targeting different marker genes and regions used for amplification of multiple fragments (amplicons) used in the Fluidigm Access Array 48 x 48 (Fluidigm, San Francisco, California, USA), for the samples from the wild-derived and wild-caught mice.

**Additional file 2:** Word document containing supplementary information: 1) Methods for redefining taxonomic annotation of ASVs annotated as “Oxyurida” at the genus level; 2) Table S1-2. Factors shaping the intestinal microbiome of wild mice. 3) Table S3. Bacteria composition similarity is predicted by fungi composition similarity in the intestinal community of wild mice. 4) Table S4. Factors shaping the intestinal microbiome of lab (inbred) mice.

## Notes

https://github.com/ferreira-scm/Hybridization_spatial_HMHZ

### Competing Interest Statement

The authors have declared no competing interest.

