## Additional file 2 for "Subspecies divergence, hybridisation and the spatial environment shape phylosymbiosis in the microbiome of house mice"

**Supplementary information: Methods for redefining taxonomic annotation of ASVs annotated as “Oxyurida” at the genus level**

Following the approach previously applied to classify *Eimeria* spp. ^48^, we first evaluated the co-abundance network structure for each ASV annotated as a known parasites at the genus level (Oxyurida, Tritrichomonas, Cryptosporidium, Eimeria, Ascaridida, Spirurida, Cyclophyllidea and Trichocephalida). ASVs annotated as “Oxyurida” at the genus level were likely from 2 different taxa as they form 2 clusters, whereas the network co-occurrence network structure suggests that each genus is composed of one taxon. We then conducted phylogenetic analysis based on the 18S rRNA gene and assigned ASVs as *Syphacia* sp. or *Aspiculuris* sp. In addition, ASVs assigned at the genus level to the orders Spirurida, Trichocephalida, and Cyclophyllidae were relabelled to the genera *Mastophorus*, *Trichuris,* and *Hymenolepis*, respectively.

We constructed a phylogenetic tree based on the 18S rRNA gene, as described for *Eimeria* spp. ^48^, including all 18S rRNA ASVs annotated as Oxyurida, and 18S rRNA gene sequences from from NCBI for *Aspiculuris* (GenBank accession codes: EF464551.1, MH215350.1, KP338606.1, KY462827.1, MT755640.1, KY462828.1, MT613322.1) and *Syphacia* (GenBank accession codes: EF464553.1, EF464554.1, AB629697.1, KY462829.1, KY462826.1, OK138900.1, OK138907.1, OK138885.1, OK138886.1, OK138897.1, OK138899.1, OK138904.1, OK138908.1, EU263105.2, MT135057.1, MT135058.1) and *Enterobius vermicularis* as the outgroup (GenBank accession codes: JF934731.1, FR687850.1, AB626598.1). The selected best fitting model was GTR+F. ASVs were assigned as *Syphacia* or *Aspiculuris*.

**Table S1.** Factors shaping the intestinal microbiome of wild mice. Distance-based models for each component of the intestinal microbiome composition among pairs of individuals (n=191,271) for occurrence-based (Jaccard similarity distances). Shown are the mean estimates of the posterior distribution for each parameter and the associated 95% credible intervals (95% CI) and R-hat value that provides information on the chain convergence (if considerably greater than 1.01, the parameter estimate is not reliable). Bold highlights significant effects (credible intervals do not overlap 0).

|  | Parasites | | | Fungi | | | Plants | | | Bacteria | | |
| --- | --- | --- | --- | --- | --- | --- | --- | --- | --- | --- | --- | --- |
|  | **Estimate** | **CI 95%** | **Rhat** | **Estimate** | **CI 95%** | **Rhat** | **Estimate** | **CI 95%** | **Rhat** | **Estimate** | **CI 95%** | **Rhat** |
| Intercept | -0.435 | -0.503: -0.365 | 1.01 | -0.723 | -0.850: -0.592 | 1.00 | -0.968 | -1.060: -0.882 | 1.01 | -1.466 | -1.549: -1.392 | 1.42 |
| Spatial distances | -0.024 | -0.058: 0.010 | 1.00 | **-0.052** | **-0.102: -0.002** | **1.00** | **-0.149** | **-0.183: -0.115** | 1.00 | **-0.067** | **-0.077: -0.057** | **1.00** |
| Subspecies’ genetic distances | **-0.047** | **-0.075: -0.018** | **1.00** | **-0.128** | **-0.167: -0.099** | **1.00** | 0.012 | -0.013: 0.037 | 1.00 | **-0.016** | **-0.024: -0.008** | **1.00** |
| hHe-dist | 0.005 | -0.035: 0.044 | 1.00 | **-0.110** | **-0.166: -0.054** | **1.00** | 0.027 | -0.010: 0.065 | 1.00 | 0.004 | -0.008: 0.015 | 1.01 |
| hHe-mean | -0.026 | -0.137: 0.087 | 1.01 | **-0.273** | **-0.493: -0.058** | **1.01** | -0.143 | -0.284: 0.015 | 1.01 | 0.040 | -0.095: 0.190 | 1.43 |
| Temporal distances | -0.012 | -0.027: 0.003 | 1.00 | **-0.046** | **-0.067: -0.026** | **1.00** | **-0.499** | **-0.513: -0.486** | 1.00 | **-0.031** | **-0.035: -0.027** | **1.00** |
| Subspecies’ genetic distances: hHe-dist | -0.016 | -0.096: 0.064 | 1.00 | 0.106 | -0.004: 0.217 | 1.00 | -0.029 | -0.102: 0.044 | 1.00 | -0.003 | -0.026: 0.020 | 1.01 |

**Table S2.**  Factors shaping the intestinal microbiome of wild mice. Distance-based models for each component of the intestinal microbiome composition among pairs of individuals (n=191,271) for abundance-based (Aitchison similarity distances). Shown are the mean estimates of the posterior distribution for each parameter and the associated 95% credible intervals (95% CI) and R-hat value that provides information on the chain convergence (if considerably greater than 1.01, the parameter estimate is not reliable). Bold highlights significant effects (credible intervals do not overlap 0).

|  | Parasites | | | Fungi | | | Plants | | | Bacteria | | |
| --- | --- | --- | --- | --- | --- | --- | --- | --- | --- | --- | --- | --- |
|  | **Estimate** | **CI 95%** | **Rhat** | **Estimate** | **CI 95%** | **Rhat** | **Estimate** | **CI 95%** | **Rhat** | **Estimate** | **CI 95%** | **Rhat** |
| Intercept | -0.061 | -0.190: 0.083 | 1.23 | -0.254 | -0.377: -0.118 | 1.14 | 0.460 | 0.383: 0.565 | 1.30 | -0.157 | -0.202: -0.106 | 1.11 |
| Spatial distances | **-0.024** | **-0.037: -0.010** | **1.00** | -0.011 | -0.031: 0.010 | 1.00 | **0.013** | **0.006: 0.019** | **1.00** | **-0.017** | **-0.021: -0.012** | **1.00** |
| Subspecies’ genetic distances | **-0.033** | **-0.044: -0.023** | **1.00** | **-0.053** | **-0.069: -0.037** | **1.00** | 0.002 | -0.002: 0.007 | 1.01 | **-0.004** | **-0.008: -0.001** | **1.00** |
| hHe-dist | 0.012 | -0.004: 0.027 | 1.01 | **-0.055** | **-0.078: -0.031** | **1.00** | 0.004 | -0.004: 0.011 | 1.01 | 0.000 | -0.004: 0.005 | 1.00 |
| hHe-mean | -0.087 | -0.326: 0.107 | 1.17 | 0.136 | -0.077: 0.352 | 1.10 | 0.047 | -0.054: 0.173 | 1.34 | -0.006 | -0.109: 0.067 | 1.08 |
| Temporal distances | **-0.011** | **-0.017: -0.006** | **1.00** | -0.005 | -0.014: 0.004 | 1.00 | **-0.107** | **-0.110: -0.104** | **1.00** | **-0.018** | **-0.020: -0.017** | **1.00** |
| Subspecies’ genetic distances: hHe-dist | -0.005 | -0.036: 0.026 | 1.01 | **0.096** | **0.049: 0.144** | **1.00** | -0.005 | -0.020: 0.009 | 1.01 | 0.001 | -0.009: 0.011 | 1.00 |

**Table S3.** Bacteria composition similarity is predicted by fungi composition similarity in the intestinal community of wild mice. Distance-based models for the overall and each component of the intestinal microbiome composition among pairs of individuals for occurrence-based (Jaccard similarity distances) and abundance-based (Aitchison similarity distances). Shown are the mean estimates of the posterior distribution for each parameter and the associated 95% credible intervals (95% CI) and R-hat value that provides information on the chain convergence (if considerably greater than 1.01, the parameter estimate is not reliable). Bold highlights significant effects (credible intervals do not overlap 0).

|  | Jaccard similarity distances | | | Aitchison similarity distances | | |
| --- | --- | --- | --- | --- | --- | --- |
|  | Estimate | CI 95% | Rhat | Estimate | CI 95% | Rhat |
| Intercept | 0.198 | 0.182: 0.212 | 1.21 | -0.143 | -0.187: -0.088 | 1.19 |
| Fungal community similarity | 0.008 | 0.007: 0.009 | 1.00 | 0.027 | 0.026: 0.028 | 1.00 |
| Spatial distances | -0.011 | -0.012: -0.009 | 1.00 | -0.016 | -0.021: -0.012 | 1.00 |
| Subspecies’ genetic distances | -0.003 | -0.004: -0.002 | 1.00 | -0.003 | -0.006: 0.001 | 1.00 |
| hHe-dist | 0.001 | -0.001: 0.003 | 1.00 | 0.002 | -0.003: 0.007 | 1.00 |
| hHe-mean | 0.008 | -0.017: 0.033 | 1.09 | -0.025 | -0.128: 0.062 | 1.23 |
| Temporal distances | -0.005 | -0.006: -0.004 | 1.00 | -0.018 | -0.020: -0.016 | 1.00 |
| Subspecies’ genetic distances: hHe-dist | -0.001 | -0.005: 0.003 | 1.00 | -0.001 | -0.011: 0.009 | 1.00 |

**Table S4.** Factors shaping the intestinal microbiome of lab (inbred) mice. Distance-based models for the overall and each component of the intestinal microbiome composition among pairs of individuals for occurrence-based (Jaccard similarity distances) and abundance-based (Aitchison similarity distances). Shown are the mean estimates of the posterior distribution for each parameter and the associated 95% credible intervals (95% CI) and R-hat value that provides information on the chain convergence (if considerably greater than 1.01, the parameter estimate is not reliable). Bold highlights significant effects (credible intervals do not overlap 0). 22 mice before and at the peak of *Eimeria* infection resulted in n= 842 dyadic comparisons (excluding self-comparisons).

| Jaccard similarity | | | | | | | | | | | | | | | |
| --- | --- | --- | --- | --- | --- | --- | --- | --- | --- | --- | --- | --- | --- | --- | --- |
|  | Overall | | | Parasites | | | Fungi | | | Plants | | | Bacteria | | |
|  | Estimate | CI 95% | Rhat | Estimate | CI 95% | Rhat | Estimate | CI 95% | Rhat | Estimate | CI 95% | Rhat | Estimate | CI 95% | Rhat |
| Intercept | **-0.917** | **-1.203: -0.636** | **1.00** | 0.142 | -0.085: 0.376 | 1.00 | **-0.340** | **-0.623: -0.050** | **1.01** | -0.264 | -0.590: 0.054 | 1.01 | **-1.098** | **-1.306: -0.888** | **1.00** |
| Subspecies’  genetic distances | **-0.085** | **-0.159: -0.010** | **1.00** | 0.070 | -0.057: 0.197 | 1.00 | 0.036 | -0.100: 0.169 | 1.00 | 0.003 | -0.127: 0.133 | 1.00 | -0.084 | -0.175: 0.006 | 1.00 |
| hHe-dist | **-0.091** | **-0.181: -0.00005** | **1.00** | 0.208 | -0.106: 0.518 | 1.00 | **-0.281** | **-0.431: -0.130** | **1.00** | -0.015 | -0.152: 0.126 | 1.00 | -0.062 | -0.182: 0.063 | 1.00 |
| Infection | **-0.329** | **-0.386: -0.271** | **1.00** | **-1.038** | **-1.161: -0.917** | **1.00** | -0.071 | -0.173: 0.031 | 1.00 | **-0.186** | **-0.284: -0.086** | **1.00** | **-0.246** | **-0.319: -0.172** | **1.00** |
| Aitchison similarity | | | | | | | | | | | | | | | |
|  | Estimate | CI 95% | Rhat | Estimate | CI 95% | Rhat | Estimate | CI 95% | Rhat | Estimate | CI 95% | Rhat | Estimate | CI 95% | Rhat |
| Intercept | 0.092 | -0.024: 0.211 | 1.00 | **0.803** | **0.750: 0.857** | **1.00** | **0.843** | **0.819: 0.867** | **1.01** | **0.666** | **0.606: 0.726** | **1.01** | **0.706** | **0.654: 0.759** | **1.01** |
| Subspecies’ genetic distances | **-0.102** | **-0.137: -0.067** | **1.00** | -0.001 | -0.031: 0.028 | 1.00 | 0.004 | -0.009: 0.017 | 1.00 | 0.002 | -0.022: 0.025 | 1.00 | -0.005 | -0.022: 0.013 | 1.00 |
| hHe-dist | 0.007 | -0.032: 0.045 | 1.00 | **-0.040** | **-0.073: -0.007** | **1.00** | -0.013 | -0.027: 0.0002 | 1.00 | -0.018 | -0.044: 0.008 | 1.00 | -0.014 | -0.032: 0.004 | 1.00 |
| Infection | **-0.166** | **-0.193: -0.139** | **1.00** | **-0.241** | **-0.264: -0.219** | **1.00** | -0.009 | -0.019: 0.0003 | 1.00 | **-0.062** | **-0.080: -0.043** | **1.00** | -0.012 | -0.024: 0.002 | 1.00 |
